# FP8 Inference in Genomic Foundation Models: Theoretical vs. Realized Speedups on GenomeOcean

**DOI:** 10.64898/2026.08.09.743676

**Authors:** Mutian Yu, Rob Egan, Fengchen Liu, Zhong Wang, Lizhen Shi

## Abstract

Genomic Foundation Models (GFMs) are increasingly used for large-scale sequence analysis and generation. Compared with frontier language models, GFMs are typically smaller and frequently operate on long genomic sequences, with evaluation often requiring preservation of biologically meaningful structure and sequence-level relationships. Although low-precision post-training quantization (PTQ) has shown substantial memory and throughput benefits for general-purpose language models, it remains unclear whether these benefits transfer to GFMs given their distinct model scales, sequence characteristics, and evaluation requirements.

We present an empirical case study of FP8 post-training quantization applied to GenomeOcean, a computationally efficient genomic foundation model with strong reported performance across diverse genomics tasks [Zhou et al., 2025]. Its range of model scales, from 100M to 4B parameters, provides a useful setting for examining how quantization effects vary with model size. We evaluate FP8 across two primary GFM inference regimes—embedding extraction and autoregressive generation—and assess its impact along two dimensions: biological fidelity relative to BF16 baselines and system-level efficiency in terms of throughput, memory usage, and energy efficiency.

We find that FP8 largely preserves biological fidelity across the evaluated scales and inference regimes, while reducing GPU memory footprint at 4B scale and improving energy efficiency during autoregressive generation. However, realized throughput gains remain substantially below FP8’s theoretical 2× hardware ceiling, with a best-case improvement of 19.3% in autoregressive generation and benefits varying strongly by model scale and workload. Autoregressive generation shows the clearest gains, driven largely by KV-cache compression, whereas embedding extraction provides limited or negative throughput benefits at smaller model scales. We attribute this theory–practice gap to the interaction of model-scale effects, memory-system bottlenecks, and software-stack limitations. These findings highlight the need for workload-specific empirical evaluation before adopting low-precision inference in scientific foundation models.

**Code availability:** https://github.com/jgi-genomeocean/genomeocean_efficiency

## 1. Introduction

The success of large language models (LLMs) has inspired the development of genomic foundation models (GFMs), which are pretrained on large-scale DNA sequence corpora using self-supervised learning. Models such as GenomeOcean [Zhou et al., 2025], Evo [Nguyen et al., 2024], and DNABERT-2 [Zhou et al., 2024] have demonstrated strong performance across diverse genomics tasks, including metagenomic species identification, regulatory sequence analysis, and functional sequence generation. As GFMs grow in capability and adoption, efficient inference and deployment become increasingly important.

Although GFMs are generally smaller than frontier NLP models — typically hundreds of millions to a few billion parameters — they still impose substantial memory and compute demands at inference. A 500M-parameter model in BF16 requires approximately 1 GB for weights alone, before accounting for activations, intermediate states, and the KV cache during autoregressive generation. At multi-billion-parameter scale, these demands can approach or exceed a single GPU’s memory, especially with long contexts or large batches. This burden is compounded by genomics workloads, which often require processing vast numbers of sequences for embedding extraction or generation. Consequently, inference cost — not merely model size — frequently becomes the binding constraint on deployment.

Post-training quantization (PTQ) is an attractive approach for reducing inference cost because it can lower memory usage and improve throughput without retraining. In large NLP models, methods such as GPTQ [Frantar et al., 2023], SmoothQuant [Xiao et al., 2023], AWQ [Lin et al., 2024], and KV-cache compression schemes [Hooper et al., 2024] have demonstrated substantial memory savings with limited accuracy degradation. However, the practical value of PTQ for GFMs remains largely unexplored. GFM inference commonly includes workloads such as large-scale embedding extraction and autoregressive sequence generation, where quantization-induced numerical perturbations may alter biologically meaningful representation geometry or sequence likelihoods. Evaluating PTQ for GFMs therefore requires considering not only system-level efficiency, but also preservation of biological fidelity.

We focus on FP8 because it combines floating-point dynamic range with native acceleration on recent GPU hardware. Compared with fixed-point formats such as INT8 [Dettmers et al., 2022], FP8’s floating-point exponent provides greater flexibility for representing heterogeneous activation magnitudes. Our preliminary activation-distribution analysis (Appendix Figure 4) further shows FP8 E4M3 more closely preserves the BF16 activation distribution of a representative GenomeOcean-4B layer than INT8 or FP4. FP8 is also natively accelerated on NVIDIA H100 Tensor Cores [NVIDIA Corporation, 2022] and supported by inference frameworks including PyTorch and vLLM [Kwon et al., 2023], making it well suited for our PTQ evaluation.

We selected GenomeOcean as our evaluation target because it is a computationally efficient genomic foundation model available at multiple model scales. Zhou et al. [Zhou et al., 2025] report up to 80× faster inference than models of comparable size for genome generation, attributed to its optimized architecture and efficient attention mechanisms. GenomeOcean therefore provides a useful setting for testing whether FP8 can deliver additional efficiency gains on top of an already optimized inference stack. Its 100M, 500M, and 4B variants also allow us to examine how quantization behavior changes with model scale.

We evaluate FP8 across two representative GFM inference regimes. For embedding extraction, we benchmark the 100M, 500M, and 4B GenomeOcean variants using PyTorch with FBGEMM FP8 kernels. For autoregressive sequence generation, we evaluate the 4B model using vLLM with FP8 KV-cache and W8A8 dynamic quantization. These two regimes expose different compute and memory bottlenecks. Embedding extraction is dominated by full-sequence forward computation, whereas autoregressive generation introduces repeated decoding and increasing KV-cache memory traffic. Although our empirical results are specific to GenomeOcean and FP8, the mechanisms we examine—including quantization overhead, arithmetic intensity, and KV-cache memory pressure—are common to transformer inference and may inform deployment of other small- and medium-scale scientific foundation models.

This paper makes the following contributions:

- **Quantitative FP8 benchmarks for GFMs.** We provide a systematic evaluation of FP8 PTQ on GenomeOcean across embedding extraction and autoregressive generation, spanning model scales from 100M to 4B parameters on NVIDIA H100 hardware.
- **Biological fidelity is largely preserved under FP8.** DBSCAN-based ARI for metagenomic binning and perplexity on GTDB Bacteria sequences remain close to BF16 baselines across the evaluated settings.
- **Realized FP8 gains are workload- and scale-dependent.** Despite an idealized 2× dense-matrix throughput advantage, the best observed end-to-end gain is 19.3% over BF16. We show that realized benefits depend strongly on model scale, workload structure, and memory pressure.

## 2. Methods and Experimental Setup

We evaluate FP8 post-training quantization on GenomeOcean using BF16 as the reference precision.^1^ This section describes the model configurations, hardware, inference protocols, datasets, and evaluation metrics.

### 2.1 Model Architectures

We evaluate three GenomeOcean model scales—100M, 500M, and 4B parameters—to examine how FP8 inference behavior varies with model scale. The models differ in hidden dimension, feed-forward width, number of transformer layers, attention-head configuration, and maximum context length.

Table 1 summarizes the architecture parameters most relevant to inference cost, together with derived estimates of feed-forward compute and KV-cache memory requirements. The KV-cache size per token is computed as 2*n*_layers_*n*_kv_*d*_head_*b*, where *b* is the number of bytes per cached element.

**Table 1:** Inference-relevant architecture parameters for the three GenomeOcean model scales, together with per-token KV-cache requirements under BF16 and FP8.

| Parameter | 100M | 500M | 4B |
| --- | --- | --- | --- |
| Hidden size $d$ | 768 | 1536 | 3072 |
| Feed-forward size $d_{\text{ff}}$ | 3072 | 6144 | 16384 |
| Number of layers $n_{\text{layers}}$ | 12 | 14 | 24 |
| Query attention heads $n_{\text{heads}}$ | 8 | 8 | 12 |
| KV heads $n_{\text{kv}}$ | 8 | 8 | 4 |
| Head dimension $d_{\text{head}}$ | 96 | 192 | 256 |
| Maximum context length $L_{\text{max}}$ | 1,024 | 1,024 | 10,240 |
| KV-cache elements per token $2n_{\text{layers}}n_{\text{kv}}d_{\text{head}}$ | 18,432 | 43,008 | 49,152 |
| KV cache per token, BF16 | 36 KB | 84 KB | 96 KB |
| KV cache per token, FP8 | 18 KB | 42 KB | 48 KB |

### 2.2 Hardware Configuration

All experiments were conducted on a single NVIDIA H100 SXM5 GPU with 80 GB of HBM3 memory and 3.35 TB/s of memory bandwidth. The H100 provides native FP8 acceleration through Hopper-generation Tensor Cores, with an idealized dense-matrix throughput of approximately 2× that of BF16.

GPU power draw was monitored using the NVIDIA Management Library (NVML) [NVIDIA Corporation, 2024] at 100 ms intervals throughout each inference workload. We report mean steady-state power draw and compute energy efficiency as throughput per unit power, as defined in Section 2.6.

### 2.3 Inference Regimes

We evaluate two inference protocols representative of GenomeOcean deployment (Figure 1). Embedding extraction performs compute-intensive full-sequence processing, whereas autoregressive generation repeatedly accesses and extends a growing key-value (KV) cache during decoding. Consequently, the benefit of reducing numerical precision may depend not only on model size, but also on whether inference is compute-bound, memory-capacity-bound, or memory-bandwidth-bound.

**Figure 1:**
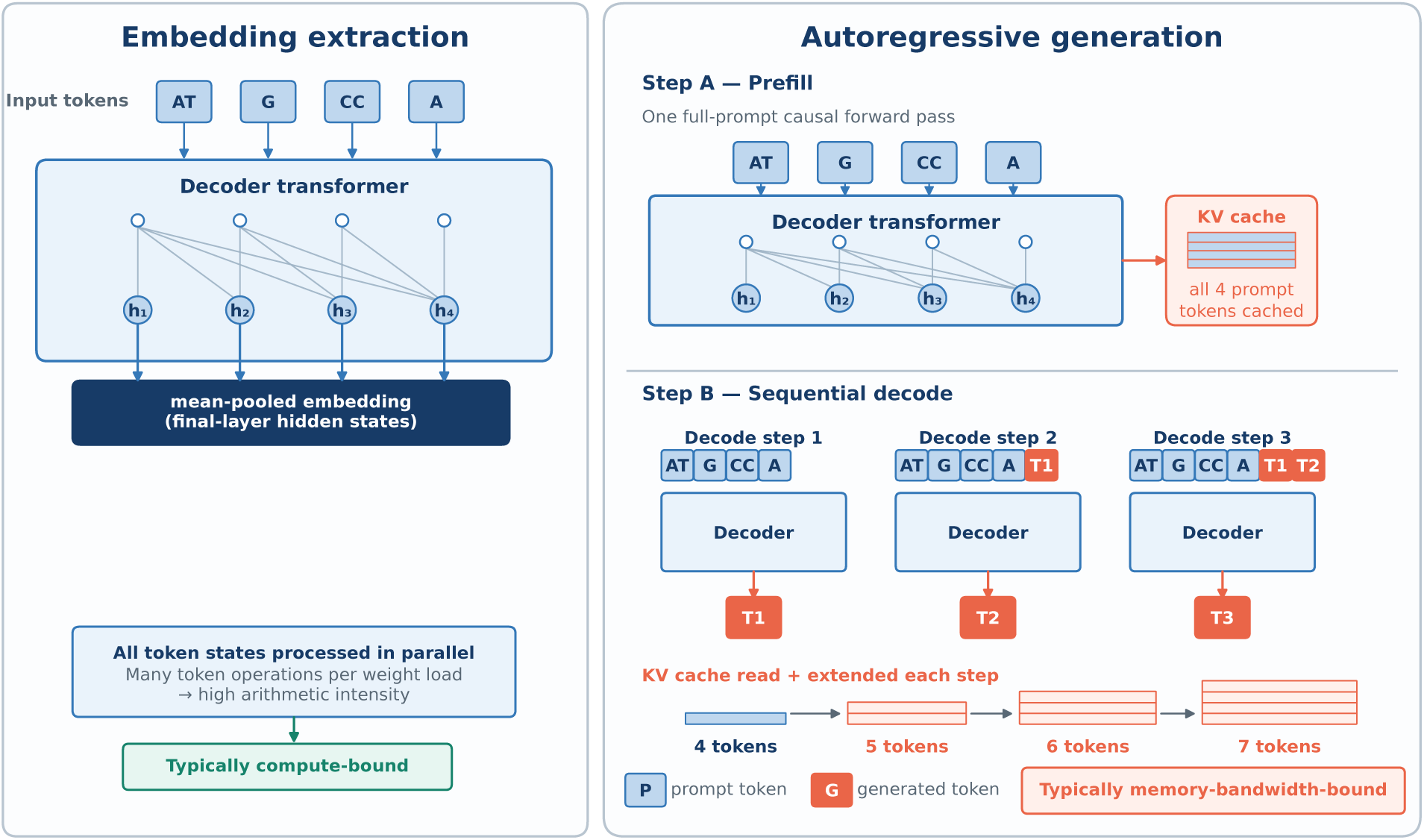
Overview of the two inference regimes evaluated in this study. **Left:** Embedding extraction performs a single full-sequence causal forward pass and returns hidden-state representations that can be pooled into sequence-level embeddings for downstream analysis. This regime is dominated by dense matrix multiplications over the full input sequence. **Right:** Autoregressive generation consists of a prefill phase that computes the KV cache, followed by incremental decoding in which each generated token repeatedly reads cached key-value states. As context length and batch size increase, decoding becomes increasingly constrained by KV-cache memory traffic [Kwon et al., 2023].

#### Embedding Extraction

The model performs a single causal forward pass over batched tokenized sequences and returns final-layer hidden states, which are mean-pooled to produce sequence-level embeddings for downstream analysis. No autoregressive decoding is performed. Throughput is measured in input tokens per second using a fixed batch size within each model scale, with the same batch size used for BF16 and FP8.

#### Autoregressive Sequence Generation

GenomeOcean-4B generates DNA sequences token by token using vLLM. Generation consists of a prefill phase that processes the input sequence and constructs the KV cache, followed by autoregressive decoding in which the cache is reused and extended for each generated token. Throughput is measured in generated tokens per second.

### 2.4 Quantization Configurations

We compare BF16 with three FP8 post-training quantization configurations (Table 2). BF16 serves as the reference precision, and all FP8 configurations use the E4M3 format without quantization-aware training or calibration fine-tuning.

**Table 2:** Summary of quantization configurations evaluated in this study. W = weights, A = activations, and KV = key-value cache. All FP8 configurations use the E4M3 format.

| Protocol | Configuration | Target | Scaling | Engine | Regime |
| --- | --- | --- | --- | --- | --- |
| A | FP8 Weight Quantization | W | Static | FBGEMM | Embedding |
| B.1 | FP8 KV Cache | KV | Dynamic (per-tensor) | vLLM | Generation |
| B.2 | W8A8 + FP8 KV Cache | W, A, KV | Static W; dynamic A/KV | vLLM | Generation |

**Protocol A** applies static FP8 weight quantization using PyTorch with FBGEMM kernels, while output embeddings remain in BF16.

For autoregressive generation, **Protocol B.1** quantizes only the KV cache to FP8, leaving weights and activations in BF16. **Protocol B.2** retains the FP8 KV cache and additionally applies FP8 quantization to weights and activations. Weights are quantized statically, whereas activations and the KV cache are scaled dynamically at runtime. The `lm head` remains in BF16 due to a vLLM implementation constraint.

Because the two inference regimes use workload-specific inference engines, absolute throughput values across Protocol A and Protocol B should not be interpreted as a controlled cross-engine comparison.

### 2.5 Datasets and Biological Fidelity Evaluation

We use separate GTDB-derived datasets to evaluate biological fidelity for embedding extraction and autoregressive generation.

#### Metagenomic binning (Protocol A)

For embedding evaluation, we construct a benchmark from the Genome Taxonomy Database (GTDB) [Parks et al., 2018]. Each trial samples 20 bacterial genomes from a pre-sampled pool of 50 genomes, with sequence fragments assigned their corresponding species labels. A representative 20-genome sample from this pool is shown in Table 5.

To maintain comparable data volume across model scales, each genome contributes approximately one million tokens. Sequences are divided into non-overlapping fragments matched to each model’s context length: 5,000-bp fragments for GenomeOcean-100M and -500M (*≈*1,024 tokens) and 50,000-bp fragments for GenomeOcean-4B (*≈*10,240 tokens).

Embedding quality is evaluated following the GenomeOcean metagenomic-binning protocol [Zhou et al., 2025]. Sequence-level embeddings are clustered using DBSCAN [Ester et al., 1996], with hyperparameters tuned on the BF16 baseline, and compared with ground-truth species labels using the Adjusted Rand Index (ARI) [Hubert & Arabie, 1985]. We report

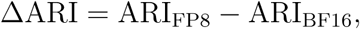

where values near zero indicate preservation of the clustering structure under quantization. This benchmark is intended as a controlled precision comparison rather than a comprehensive evaluation of taxonomic classification.

#### GTDB Bacteria sequence likelihood (Protocol B)

For autoregressive-generation fidelity, we sample 96 microbial genome sequences from GTDB Bacteria. Each genome is bounded to 102,400 tokens, and perplexity is computed using a sliding window with a 10,240-token context length and a 2,560-token stride. At each window, only the newly evaluated 2,560 tokens contribute to the negative log-likelihood, avoiding repeated evaluation of overlapping prefix tokens. Per-genome perplexity is averaged across windows to obtain one value per sequence.

We report

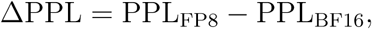

where values near zero indicate that quantization minimally changes the model’s predictive distribution over the evaluated genomic sequences.

### 2.6 System-Level Efficiency Metrics

We evaluate deployment efficiency using throughput, peak GPU memory consumption, and power efficiency.

#### Throughput

Inference throughput is measured in input tokens per second for embedding extraction and generated tokens per second for autoregressive generation. Relative change with respect to the BF16 baseline is computed as

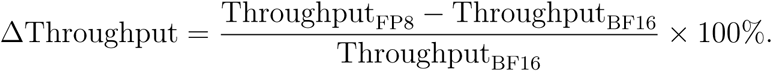

#### GPU memory consumption

Peak GPU memory allocation is reported in gigabytes. For embedding extraction, memory usage is measured using `torch.cuda.max memory allocated()`, whereas generation memory usage is obtained from vLLM’s internal memory profiler.

#### Power efficiency

GPU power draw is sampled continuously using NVML at 100 ms intervals. Operational efficiency is reported as tokens per watt (tok/W), computed as throughput divided by mean GPU power draw.

### 2.7 Experimental Repetition and Reporting

All benchmark configurations were repeated across five trials using random seeds 42–46. For Protocol A, the random seed controls genome and fragment subsampling as well as stochastic UMAP initialization; variation in the UMAP projection can in turn affect the subsequent DBSCAN clustering. For Protocol B, throughput, power, and efficiency are reported across repeated benchmark trials, whereas perplexity is computed on a fixed set of 96 GTDB Bacteria sequences. Accordingly, the reported perplexity standard deviation reflects sequence-level heterogeneity rather than trial-to-trial resampling variability.

## 3. Results

We first evaluate FP8 for embedding extraction across three GenomeOcean model scales, then examine autoregressive generation with the 4B model. Finally, we analyze time per output token across batch sizes and context lengths to characterize when FP8 KV-cache compression provides a decoding advantage.

### 3.1 Embedding Extraction: FP8 Preserves Embedding Fidelity but Shows Scale-Dependent Throughput

FP8 weight quantization preserved embedding fidelity across all three GenomeOcean model scales, but its throughput effect varied strongly with model size (Table 3). Relative to BF16, throughput decreased by 24.9% and 11.1% for the 100M and 500M models, respectively, but increased by 4.7% for the 4B model. Peak VRAM changed little at the 100M and 500M scales, whereas FP8 reduced peak VRAM by 4.23 GB for the 4B model.

**Table 3:** Protocol A embedding extraction performance and biological fidelity. Throughput and DBSCAN ARI are reported as mean *±* standard deviation over five trials. Throughput change is computed relative to the corresponding BF16 baseline.

| Model | Precision | Throughput (tok/s) | Change | Peak VRAM (GB) | DBSCAN ARI |
| --- | --- | --- | --- | --- | --- |
| 100M | BF16 | 613,371 ± 22,439 | – | 21.15 | 0.3608 ± 0.0619 |
| 100M | FP8 (E4M3) | 460,733 ± 1,694 | –24.9% | 21.03 | 0.3533 ± 0.0588 |
| 500M | BF16 | 230,523 ± 651 | – | 47.19 | 0.3345 ± 0.0479 |
| 500M | FP8 (E4M3) | 204,973 ± 777 | –11.1% | 46.66 | 0.3273 ± 0.0508 |
| 4B | BF16 | 21,386 ± 126 | – | 72.14 | 0.5278 ± 0.0672 |
| 4B | FP8 (E4M3) | 22,401 ± 204 | +4.7% | 67.91 | 0.5280 ± 0.0653 |

BF16, improves throughput by 14.7% and energy efficiency by 19.4% relative to BF16. Protocol B.2 retains the FP8 KV cache and additionally quantizes weights and activations, increasing the total throughput gain to 19.3% and the energy-efficiency gain to 27.4%. The incremental throughput gain from B.1 to B.2 is only 4.0%.

DBSCAN clustering performance remained stable across model scales. The absolute difference in mean ARI between BF16 and FP8 was below 0.008 for all three models, with essentially no change for GenomeOcean-4B (0.5278 for BF16 versus 0.5280 for FP8). Thus, FP8 preserved the evaluated clustering signal across model sizes, while a throughput improvement was observed only for the largest model tested.

Direct comparisons between BF16 and FP8 embeddings further support the conclusion that quantization introduces only small changes in the evaluated embedding representations. Across the evaluated model scales and layers, cosine error remained at or below 0.0011 and KL divergence at or below 0.0474 (Figure 2). Supplementary clustering metrics and representative UMAP visualizations in Appendix 2 show similarly small BF16–FP8 differences.

**Figure 2:**
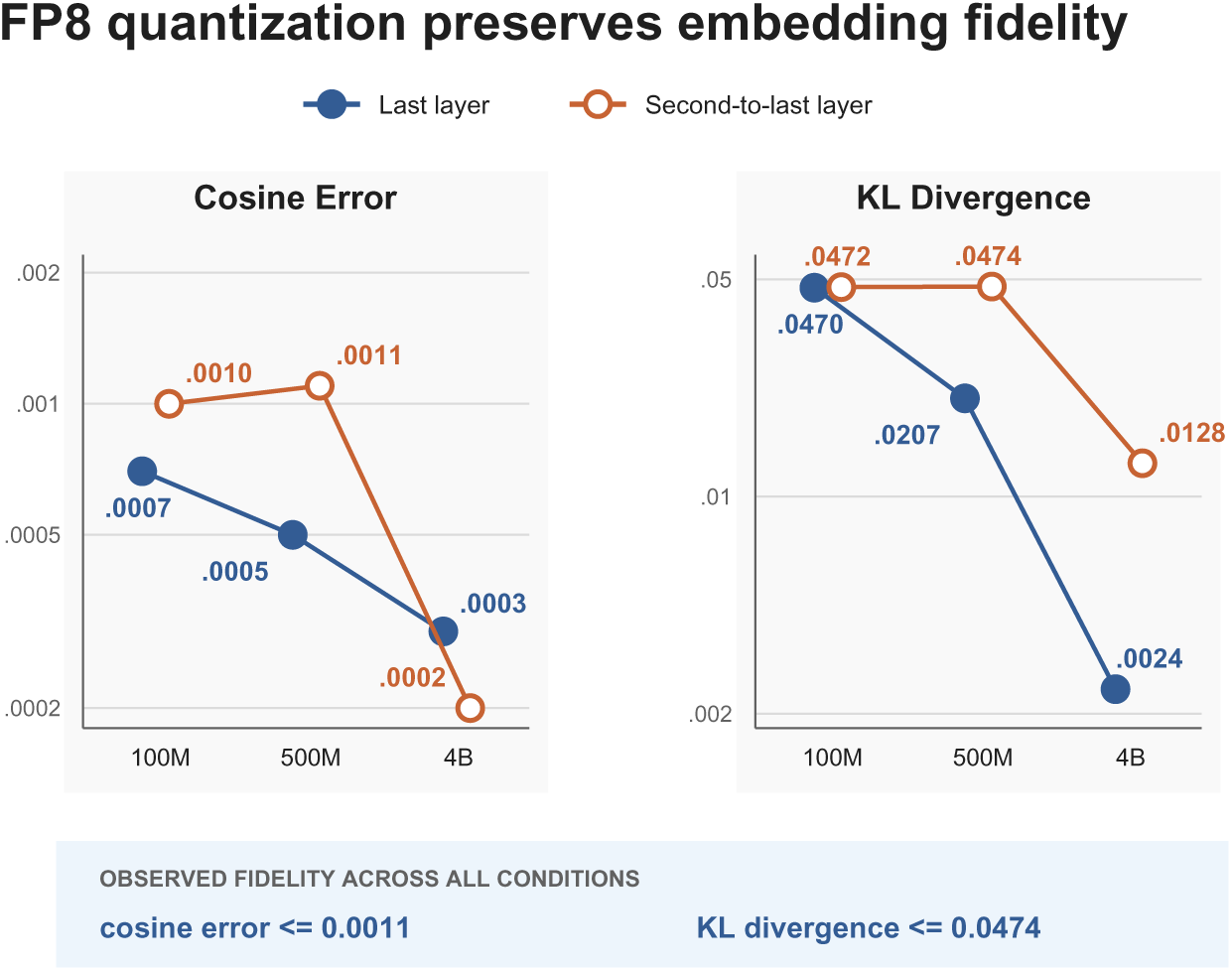
FP8 embedding fidelity metrics relative to BF16 reference embeddings for GenomeOcean-100M, -500M, and -4B. Cosine error and KL divergence show small deviations between FP8 and BF16 embeddings across the evaluated layers.

### 3.2 Autoregressive Generation: FP8 Improves Throughput While Largely Preserving Perplexity

Protocol B evaluated autoregressive sequence generation in vLLM using the GenomeOcean-4B model. As summarized in Table 4, both FP8 configurations improve throughput and energy efficiency relative to BF16.

**Table 4:** Protocol B generation benchmarks for GenomeOcean-4B. Throughput, power, and efficiency are reported as mean *±* standard deviation across 5 trials. Perplexity is reported as mean *±* standard deviation across 96 fixed GTDB Bacteria sequences, with the standard deviation reflecting sequence-level heterogeneity rather than trial-to-trial resampling variability.

| Protocol | Quantization Scheme | Throughput (tok/s) | Power (W) | Efficiency (tok/W) | Perplexity (± SD) |
| --- | --- | --- | --- | --- | --- |
| Baseline | BF16 | 1818.79 ± 7.95 | 692.40 ± 1.88 | 2.63 ± 0.01 | 125.97 ± 1.04 |
| B.1 | FP8 KV-Cache | 2085.89 ± 34.49 | 663.77 ± 8.84 | 3.14 ± 0.02 | 126.63 ± 1.04 |
| B.2 | W8A8 + FP8 KV-Cache | 2169.04 ± 24.30 | 647.01 ± 8.55 | 3.35 ± 0.02 | 128.73 ± 1.04 |

Most of the observed generation-side throughput gain is achieved through KV-cache quantization. Protocol B.1, which applies FP8 to the KV cache while leaving weights and activations in BF16, improves throughput by 14.7% and energy efficiency by 19.4% relative to BF16. Protocol B.2 retains the FP8 KV cache and additionally quantizes weights and activations, increasing the total throughput gain to 19.3% and the energy-efficiency gain to 27.4%. The incremental throughput gain from B.1 to B.2 is only 4.0%.

These efficiency gains are accompanied by only small changes in perplexity on the GTDB Bacteria sequences. Mean perplexity increases by 0.66 points under Protocol B.1 and 2.76 points under Protocol B.2 relative to BF16, corresponding to a maximum relative increase of approximately 2.2%. Because perplexity is evaluated on a fixed set of 96 GTDB Bacteria sequences, the reported standard deviation reflects sequence-level heterogeneity rather than trial-to-trial resampling variability.

### 3.3 Autoregressive Decoding: FP8 KV-Cache Helps at Large Batch and Long Context

The aggregate Protocol B results show that FP8 KV-cache compression accounts for most of the realized generation-side throughput gain. We next ask when this benefit emerges during decoding by analyzing time per output token (TPOT) during continuous autoregressive generation with GenomeOcean-4B (Figure 3). Sequences from the Protocol B generation dataset were decoded from a fixed 1,024-token prefix, followed by 9,216 generated tokens, across batch sizes 16, 32, 64, and 128. These settings span the model’s 10,240-token maximum context length. TPOT values are averaged across five independent trials and plotted as a function of effective context length.

**Figure 3:**
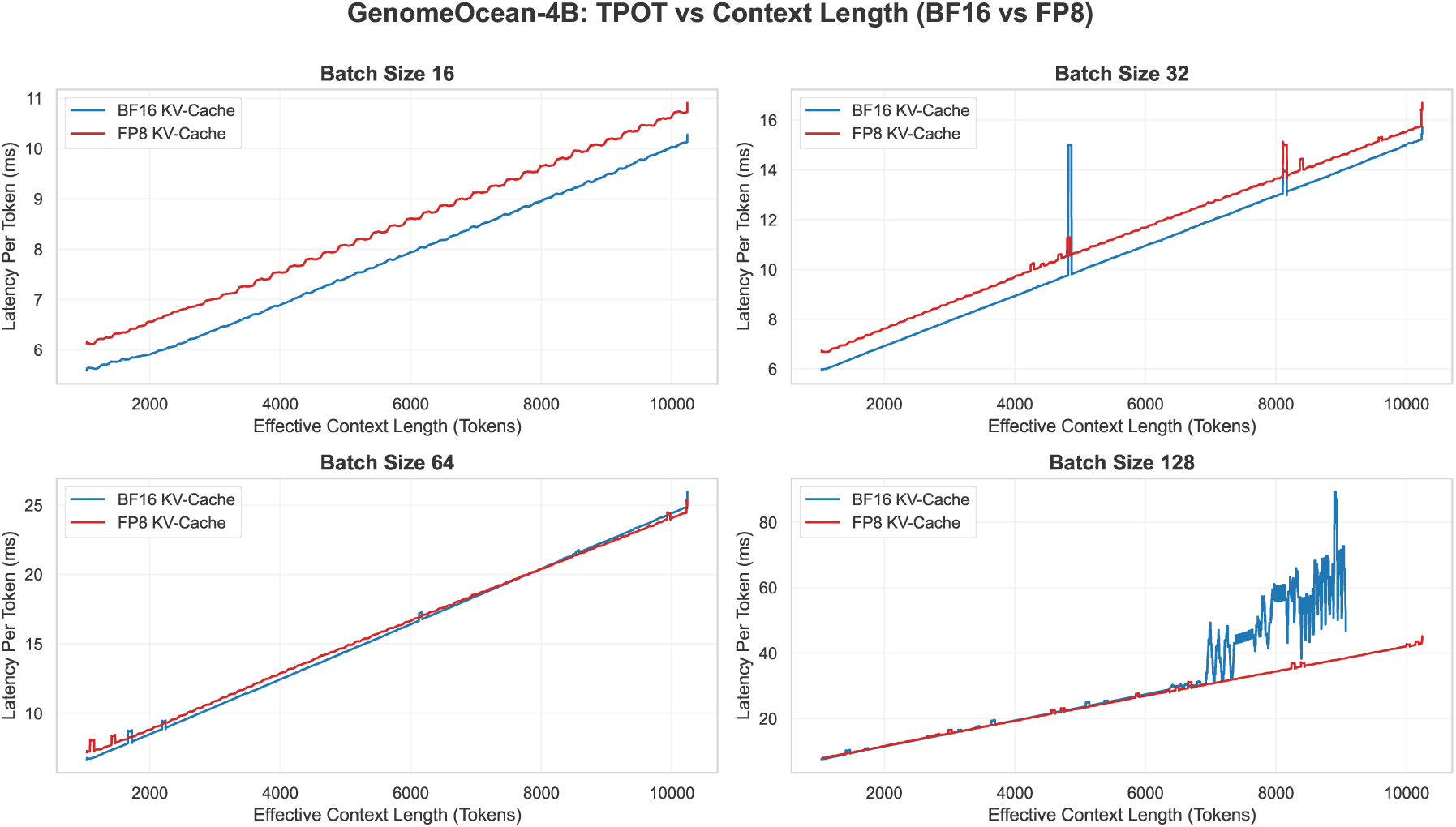
Time Per Output Token (TPOT) as a function of effective context length during continuous autoregressive generation. At smaller batch sizes (16 and 32), FP8 KV-cache overhead yields higher latency than BF16. At larger batch sizes, memory bandwidth dominates and FP8 KV-cache compression reduces latency. The BF16 batch-128 curve is truncated near 9,080 tokens, where the KV cache exceeds VRAM and vLLM triggers block-level swapping [Kwon et al., 2023].

At smaller batch sizes (16 and 32), FP8 KV-cache generation is slower than BF16, indicating that the overhead of FP8 scaling, casting, and KV-cache conversion exceeds the memory-bandwidth savings. As batch size increases, the performance gap narrows and then reverses: at batch sizes of 64 and above, KV-cache memory traffic becomes the dominant bottleneck, and the 50% reduction in KV-cache size provided by FP8 yields lower per-token latency. At batch size 128, the BF16 run exhibits severe latency fluctuations near an effective context length of approximately 9,080 tokens, where the KV cache exceeds available VRAM and vLLM triggers block-level swapping to host memory [Kwon et al., 2023]. The plotted curve is truncated at this point to avoid reporting the artificially low latency of the remaining fractured batch as if the full batch were still active.

Taken together, these results separate fidelity from efficiency: FP8 largely preserves the evaluated biological signals across GenomeOcean workloads, while its performance benefits depend on model scale, workload structure, and KV-cache memory pressure. Embedding extraction preserves clustering structure and embedding geometry under FP8, yet throughput decreases for the 100M model and improves only modestly for the 4B model. Autoregressive generation shows only small changes in perplexity while providing clearer throughput and energy-efficiency gains, particularly when FP8 is applied to the KV cache. These results indicate that FP8 should be treated as a workload-specific optimization rather than a uniformly beneficial replacement for BF16.

## 4. Discussion

FP8 post-training quantization largely preserves biological fidelity across both inference regimes but delivers realized speedups substantially below the idealized 2× dense-matrix through-put ceiling. We trace this gap to three interacting sources: model-scale effects, memory-system bottlenecks, and software-stack limitations.

### 4.1 Scale-Dependent Effects: Arithmetic Intensity and Quantization Overhead

The Protocol A results show that FP8 is not uniformly beneficial for embedding extraction. Throughput decreases by 24.9% and 11.1% relative to BF16 for the 100M and 500M models, respectively, but increases by 4.7% for the 4B model. This progression is consistent with greater amortization of FP8 quantization overhead as model and workload scale increase. In the smaller configurations, fixed costs associated with kernel dispatch, scaling metadata handling, and mixed-precision casting [Dettmers et al., 2022] may offset the computational benefit of FP8-accelerated matrix multiplication. The negligible peak-VRAM reduction at the 100M and 500M scales further suggests that non-weight memory, including activations, intermediate buffers, and runtime over-head, accounts for a substantial fraction of the measured memory footprint, limiting the benefit of weight compression alone.

At 4B parameters, FP8 begins to provide net system benefit, improving throughput by 4.7% and reducing peak VRAM by 4.23 GB. This crossover is consistent with larger workloads better amortizing the overhead associated with low-precision execution. However, the observed throughput gain remains far below the idealized 2× dense-matrix FP8 advantage because end-to-end inference includes operations that do not receive the same acceleration, including normalization, softmax, scaling, casting, and BF16 intermediate computations. Thus, increasing model and workload scale makes FP8 benefits more visible, but does not by itself translate the hardware-level FP8 advantage into a comparable end-to-end speedup.

### 4.2 Memory-System Bottlenecks: KV-Cache Compression and Workload Structure

Autoregressive generation differs from embedding extraction because inference includes both a compute-intensive prefill phase and an iterative decoding phase that repeatedly accesses and extends the KV cache. As context length and batch size increase, this repeated cache access places increasing pressure on GPU memory capacity and bandwidth [Kwon et al., 2023; Pope et al., 2023]. This explains why KV-cache compression is the dominant source of Protocol B speedup. Protocol B.1, which quantizes only the KV cache, captures most of the realized generation-side throughput gain. Protocol B.2 adds W8A8 weight and activation quantization on top of the same FP8 KV cache, but provides only a small incremental improvement, indicating that decoding is limited more by KV-cache movement than by additional weight/activation compression.

The TPOT analysis in Section 3.3 supports this mechanism. FP8 KV-cache generation is slower than BF16 at small batch sizes, where scaling and casting overhead dominate, but becomes faster once larger batches and longer contexts make KV-cache memory traffic the limiting factor. The BF16 instability at batch size 128 further illustrates this memory pressure: once the KV cache exceeds available VRAM, vLLM begins block-level swapping, fracturing the batch and sharply increasing latency variability.

### 4.3 Software-Stack Limitations: Runtime Overhead and Incomplete FP8 Execution

The small incremental gain from Protocol B.1 to B.2 suggests that the additional benefit of W8A8 quantization is partly offset by runtime overhead. Protocol B.2 uses dynamic activation scaling to accommodate input-dependent activation ranges, requiring scaling-factor computation, FP8 casts, and precision conversions during inference. These operations add per-token overhead and help explain why B.2 improves only modestly beyond the FP8 KV-cache configuration.

The remaining gap between the realized Protocol B.2 throughput gain and the theoretical 2× dense-matrix ceiling also reflects incomplete end-to-end FP8 execution. Some operations remain in higher precision, and the vLLM constraint that keeps the `lm_head` in BF16 further illustrates that the inference path is not fully FP8. More detailed kernel-level profiling would be required to quantify the contribution of individual software-stack overheads.

### 4.4 Energy Efficiency and Broader Implications

Despite modest throughput gains, Protocol B.2 improves energy efficiency by 27.4% in tokens per watt by increasing throughput while reducing average power draw. For large scientific inference workloads, this improvement may reduce operational energy use even when end-to-end speedup is limited [Patterson et al., 2021].

The mechanisms observed here arise largely from general transformer inference behavior rather than genomics-specific properties, and may therefore inform deployment of other small- and medium-scale scientific foundation models. However, the biological-fidelity metrics are benchmark-specific: ARI measures embedding preservation for one metagenomic binning task, while perplexity measures sequence-likelihood preservation on GTDB Bacteria rather than the biological functionality of generated sequences.

Our conclusions are also bounded by the experimental setting: one model family (GenomeOcean), one precision format (FP8), one hardware platform (NVIDIA H100 SXM5), and two workload-native inference stacks (PyTorch with FBGEMM and vLLM). Thus, cross-regime differences reflect not only numerical precision but also engine-specific execution and memory behavior. GenomeOcean provides a demanding case study because it is already optimized for efficient inference [Zhou et al., 2025]; results for other model architectures, quantization formats, GPUs, and inference engines require dedicated empirical evaluation.

## 5. Conclusion

We present an empirical case study of FP8 PTQ on GenomeOcean, examining the gap between theoretical and realized inference efficiency in GFMs. Biological fidelity is largely preserved across both inference regimes: DBSCAN ARI changes only minimally across model scales, and mean perplexity increases by less than 3 points under Protocol B.2 (W8A8 + FP8 KV-cache quantization). However, realized throughput gains fall substantially short of the idealized 2× FP8 dense-matrix ceiling, ranging from a 24.9% reduction for 100M embedding extraction to a 19.3% best-case gain for autoregressive generation. Our results indicate that this theory–practice gap reflects the interaction of scale-dependent quantization overhead, memory-system pressure, and software-stack limitations. TPOT analysis further shows that FP8 generation is slower than BF16 at small batch sizes but becomes advantageous as batch size and context length increase, consistent with the growing importance of KV-cache memory traffic.

The practical value of FP8 therefore depends on the deployment bottleneck rather than throughput alone. At 4B scale, FP8 reduces Protocol A peak VRAM by 4.23 GB and halves the per-token KV cache from 96 KB to 48 KB, increasing the memory available for larger batches and longer contexts. In generation, the observed 14.7–19.3% throughput gains are accompanied by a 27.4% improvement in energy efficiency under Protocol B.2. FP8 should therefore be treated as a capacity- and efficiency-oriented deployment tool whose performance benefit is workload-dependent. Practitioners should benchmark their specific architecture and workload rather than assume that theoretical or LLM-scale quantization speedups will translate directly to scientific foundation models.

## 6. Funding Statement

The work conducted by the U.S. Department of Energy Joint Genome Institute (https://ror.org/04xm1d337), a DOE Office of Science User Facility, is supported by the Office of Science of the U.S. Department of Energy operated under Contract No. DE-AC02-05CH11231.

This research used the Lawrencium computational cluster resource provided by the IT Division at the Lawrence Berkeley National Laboratory (Supported by the Director, Office of Science, Office of Basic Energy Sciences, of the U.S. Department of Energy under Contract No. DE-AC02-05CH11231)

## Author Contributions

M.Y. implemented the benchmarking and quantization workflows, conducted the experiments, analyzed the results, generated the figures, and drafted the manuscript. L.S. conceived and supervised the study, contributed to the experimental design and interpretation of the results, and revised the manuscript. Z.W., R.E., and F.L. contributed to the conceptualization of the study, provided domain expertise, and contributed to the interpretation of the results. All authors reviewed and approved the final manuscript.

## Appendix

### 1 Activation Outlier Visualizations

Figure 4 compares activation distributions from Transformer Layer 12 of GenomeOcean-4B under BF16, INT8, FP8 E4M3, and FP4 E2M1 precision. We use Layer 12 because GenomeOcean-4B contains 24 transformer layers, making it a representative intermediate layer away from both input-adjacent and final layers. The BF16 distribution serves as the reference. INT8 preserves only a coarse approximation of the dense bulk distribution and rare extreme outliers, whereas FP8 E4M3 more closely tracks both regions, yielding substantially lower quantization error (MAE = 0.00146) than INT8 (MAE = 0.06259). FP4 E2M1 provides greater nominal compression but collapses the distribution into very few representable values, increasing error (MAE = 0.06467) and requiring newer hardware support such as NVIDIA Blackwell. These observations motivate FP8 E4M3 as the practical low-precision format for this study: it balances distributional fidelity, compression, and native acceleration on H100 Tensor Cores.

**Figure 4:**
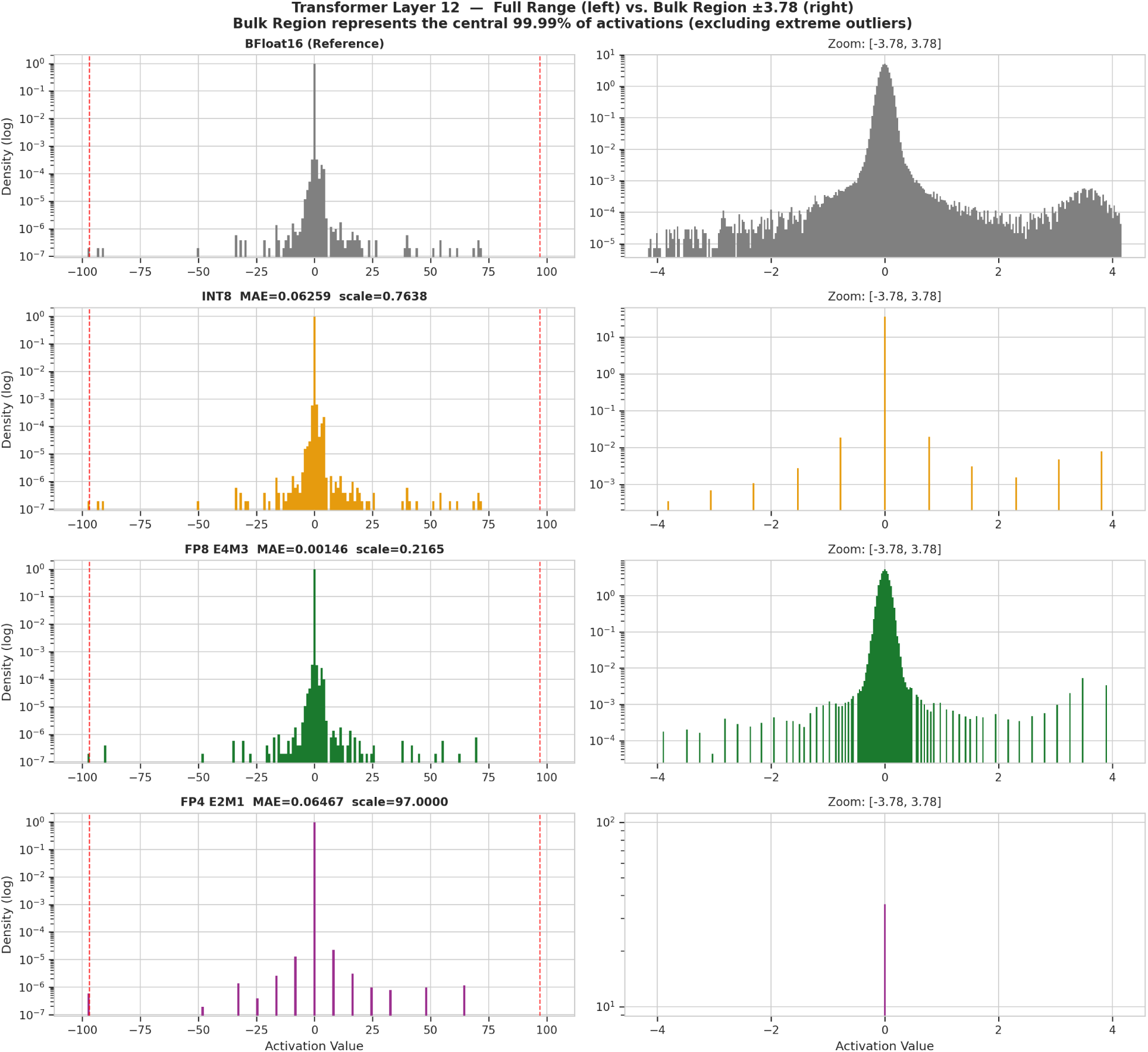
Activation distributions for Transformer Layer 12 of GenomeOcean-4B. The left panels show the full activation range, illustrating extreme outliers, while the right panels zoom in on the central 99.99% of the probability mass (*±*3.78). FP8 E4M3 (MAE = 0.00146) more closely preserves the BF16 reference distribution than INT8 (MAE = 0.06259). FP4 E2M1 (MAE = 0.06467) restricts the space to 16 values, increasing error and requiring next-generation hardware for acceleration.

### 2 Supplementary Embedding Fidelity Visualizations

Additional Protocol A visualizations support the embedding-fidelity results in Section 3.1. Figure 5 shows a representative UMAP projection, and Figure 6 reports clustering metrics beyond ARI. Both analyses show minimal BF16–FP8 differences.

**Figure 5:**
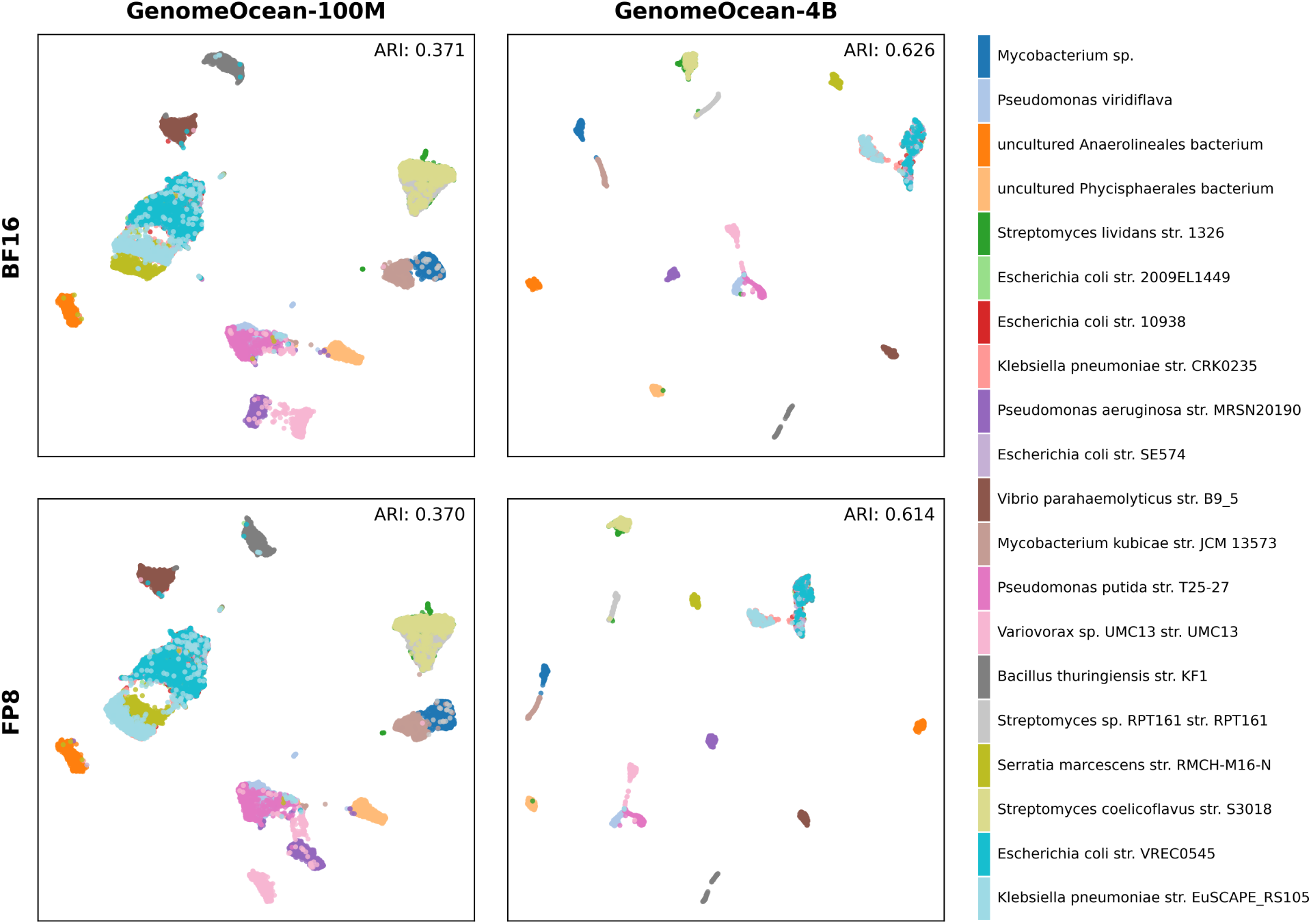
Representative UMAP visualizations of Protocol A metagenomic clustering under BF16 and FP8 precision for GenomeOcean-100M and -4B. Random state 45 is shown as an illustrative trial.

**Figure 6:**
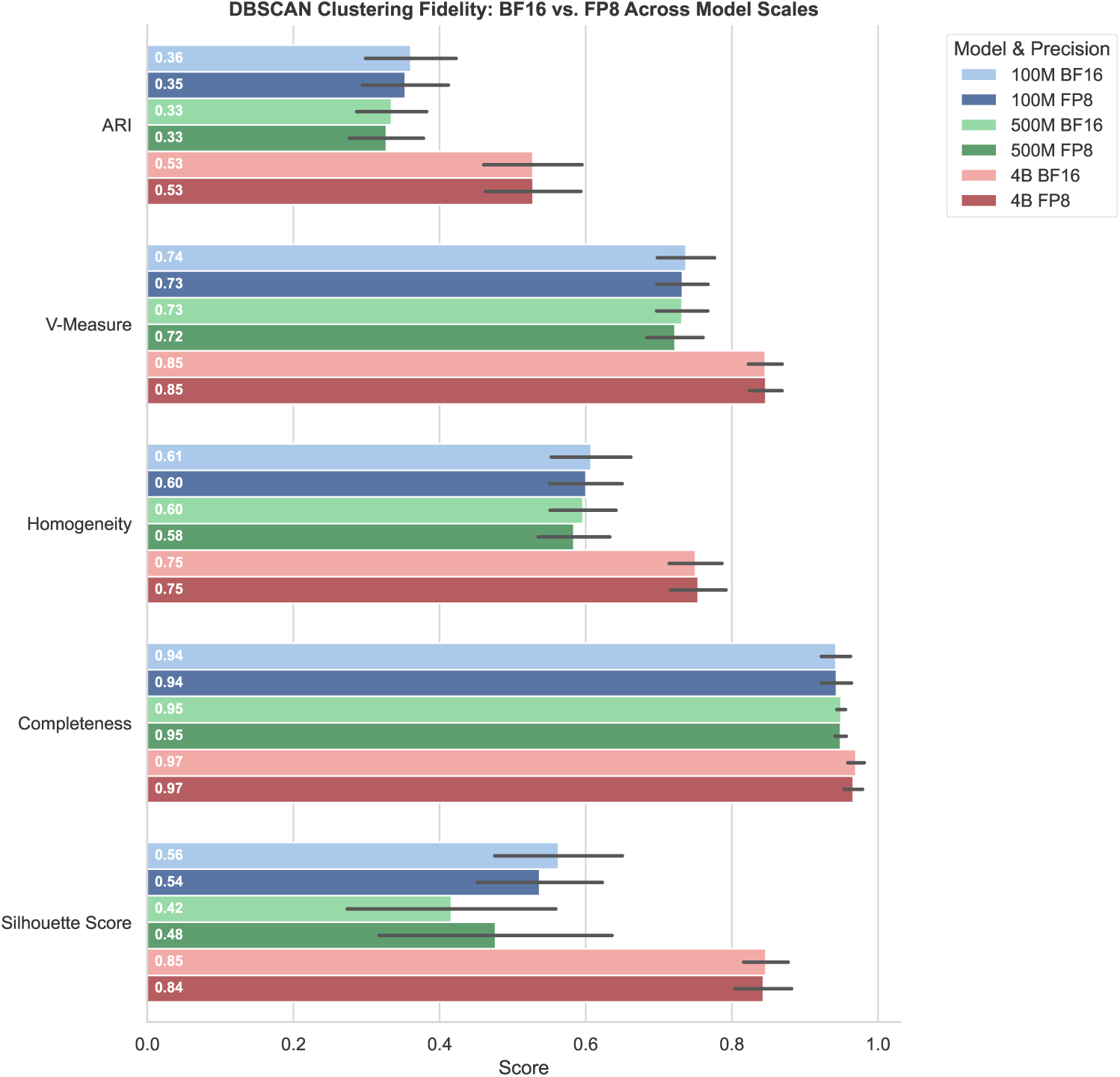
Supplementary clustering metrics comparing BF16 and FP8 embeddings across GenomeOcean-100M, -500M, and -4B. Metrics include ARI, V-measure, homogeneity, completeness, and silhouette score. Error bars represent *±*1 standard deviation across 5 trials.

### 3 Protocol A Dataset Composition

Table 5 details a representative 20-genome sample from the 50 GTDB strains utilized in Protocol A, including the GTDB accession ID, strain-level taxonomy, and the proportion of the genome sampled for evaluation. Common suffixes like “whole genome shotgun sequence” are omitted for brevity.

**Table 5:**
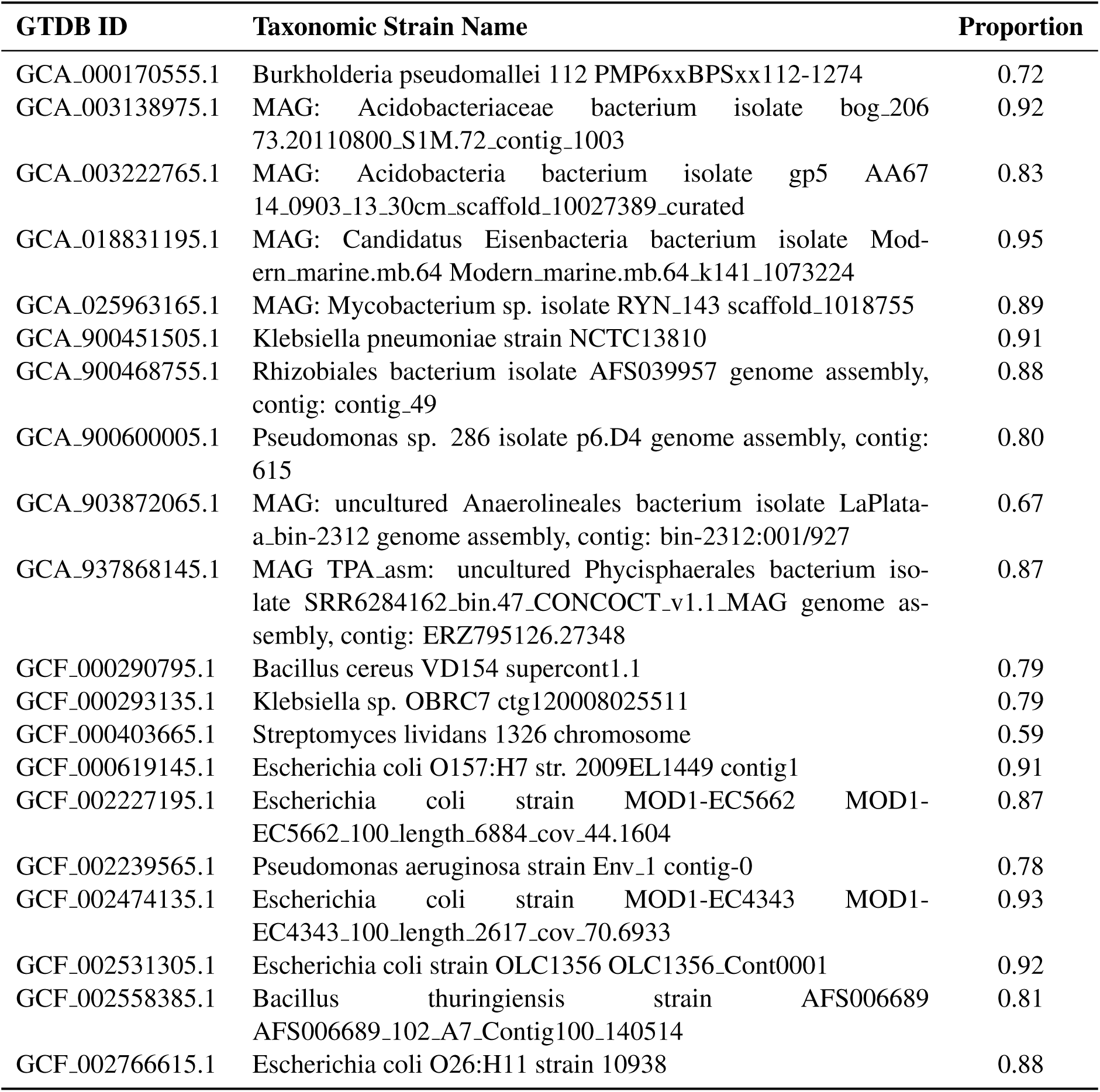
Representative 20 genomes from the Protocol A evaluation pool, detailing the GTDB ID, taxonomic strain name, and the proportion of the genome sampled for the benchmark.

The sample exhibits high intra-genus and intra-species homogeneity, containing multiple distinct strains of *Escherichia coli*, *Klebsiella pneumoniae*, and *Pseudomonas*. Because these highly similar sequences naturally overlap in the latent space, unsupervised clustering is inherently difficult. This biological homogeneity explains the relatively low absolute Adjusted Rand Index (ARI) scores (0.3–0.5) reported in Section 3.1, reinforcing that the benchmark’s goal is measuring relative precision degradation rather than perfect taxonomic separation.

## Footnotes

1 Code used for dataset construction, quantization, benchmarking, and figure generation is available at https://github.com/jgi-genomeocean/genomeocean_efficiency.git.

## Notes

https://github.com/jgi-genomeocean/genomeocean_efficiency

### Competing Interest Statement

The authors have declared no competing interest.

## References

Dettmers, T., Lewis, M., Belkada, Y., & Zettlemoyer, L. [2022]. Gpt3.int8(): 8-bit matrix multiplication for transformers at scale. In S. Koyejo, S. Mohamed, A. Agarwal, D. Belgrave, K. Cho, & A. Oh [Eds.], Advances in neural information processing systems [pp. 30318– 30332, Vol. 35]. Curran Associates, Inc. https://proceedings.neurips.cc/paper_files/paper/2022/file/c3ba4962c05c49636d4c6206a97e9c8a-Paper-Conference.pdf

Ester, M., Kriegel, H.-P., Sander, J., Xu, X., et al. [1996]. A density-based algorithm for discovering clusters in large spatial databases with noise. Kdd, 96[34], 226–231.

Frantar, E., Ashkboos, S., Hoefler, T., & Alistarh, D. [2023, March 22]. GPTQ: Accurate Post-Training Quantization for Generative Pre-trained Transformers. arXiv: 2210 . 17323 [cs]. 10.48550/arXiv.2210.17323

Hooper, C., Kim, S., Mohammadzadeh, H., Mahoney, M. W., Shao, Y. S., Keutzer, K., & Gholami, A. [2024]. KVQuant: Towards 10 million context length LLM inference with KV cache quantization. In A. Globerson, L. Mackey, D. Belgrave, A. Fan, U. Paquet, J. Tomczak, & C. Zhang [Eds.], Advances in neural information processing systems [pp. 1270–1303, Vol. 37]. Curran Associates, Inc. 10.52202/079017-0040

Hubert, L., & Arabie, P. [1985]. Comparing partitions. Journal of Classification, 2[1], 193–218. 10.1007/BF01908075

Kwon, W., Li, Z., Zhuang, S., Sheng, Y., Zheng, L., Yu, C. H., Gonzalez, J., Zhang, H., & Stoica, I. [2023]. Efficient Memory Management for Large Language Model Serving with PagedAttention. Proceedings of the 29th Symposium on Operating Systems Principles, 611–626. 10.1145/3600006.3613165

Lin, J., Tang, J., Tang, H., Yang, S., Chen, W.-M., Wang, W.-C., Xiao, G., Dang, X., Gan, C., & Han, S. [2024]. AWQ: Activation-aware weight quantization for on-device LLM compression and acceleration. In P. Gibbons, G. Pekhimenko, & C. D. Sa [Eds.], Proceedings of machine learning and systems [pp. 87–100, Vol. 6]. https://proceedings.mlsys.org/paper_files/paper/2024/file/42a452cbafa9dd64e9ba4aa95cc1ef21-Paper-Conference.pdf

Nguyen, E., Poli, M., Durrant, M. G., Kang, B., Katrekar, D., Li, D. B., Bartie, L. J., Thomas, A. W., King, S. H., Brixi, G., Sullivan, J., Ng, M. Y., Lewis, A., Lou, A., Ermon, S., Baccus, S. A., Hernandez-Boussard, T., Ré, C., Hsu, P. D., & Hie, B. L. [2024]. Sequence modeling and design from molecular to genome scale with Evo. Science, 386[6723], eado9336. 10.1126/science.ado9336

NVIDIA Corporation. [2022]. NVIDIA H100 Tensor Core GPU Architecture. white paper. https://resources.nvidia.com/en-us-hopper-architecture/nvidia-h100-tensor-c

NVIDIA Corporation. [2024]. NVIDIA management library (NVML). manual. https://developer.nvidia.com/nvidia-management-library-nvml

Parks, D. H., Chuvochina, M., Waite, D. W., Rinke, C., Skarshewski, A., Chaumeil, P.-A., & Hugenholtz, P. [2018]. A standardized bacterial taxonomy based on genome phylogeny substantially revises the tree of life. Nature Biotechnology, 36[10], 996–1004. 10.1038/nbt.4229

Patterson, D., Gonzalez, J., Le, Q., Liang, C., Munguia, L.-M., Rothchild, D., So, D., Texier, M., & Dean, J. [2021, April 23]. Carbon Emissions and Large Neural Network Training. arXiv: 2104.10350 [cs]. 10.48550/arXiv.2104.10350

Pope, R., Douglas, S., Chowdhery, A., Devlin, J., Bradbury, J., Heek, J., Xiao, K., Agrawal, S., & Dean, J. [2023]. Efficiently scaling transformer inference. In D. Song, M. Carbin, & T. Chen [Eds.], Proceedings of machine learning and systems [pp. 606–624, Vol. 5]. Curan. https://proceedings.mlsys.org/paper_files/paper/2023/file/c4be71ab8d24cdfb45e3d06dbfca2780-Paper-mlsys2023.pdf

Xiao, G., Lin, J., Seznec, M., Wu, H., Demouth, J., & Han, S. [2023, July 23–29]. SmoothQuant: Accurate and efficient post-training quantization for large language models. In A. Krause, E. Brunskill, K. Cho, B. Engelhardt, S. Sabato, & J. Scarlett [Eds.], Proceedings of the 40th international conference on machine learning [pp. 38087–38099, Vol. 202]. PMLR. https://proceedings.mlr.press/v202/xiao23c.html

Zhou, Z., Ji, Y., Li, W., Dutta, P., Davuluri, R., & Liu, H. [2024, March 18]. DNABERT-2: Efficient Foundation Model and Benchmark For Multi-Species Genome. arXiv: 2306.15006 [q-bio]. 10.48550/arXiv.2306.15006

Zhou, Z., Riley, R., Kautsar, S., Wu, W., Egan, R., Hofmeyr, S., Goldhaber-Gordon, S., Yu, M., Ho, H., Liu, F., Chen, F., Morgan-Kiss, R., Shi, L., Liu, H., & Wang, Z. [2025, February 5]. GenomeOcean: An Efficient Genome Foundation Model Trained on Large-Scale Metagenomic Assemblies. 10.1101/2025.01.30.635558

